# Pericyte ontogeny: the use of chimeras to track a cell lineage of diverse germ line origins

**DOI:** 10.1101/149922

**Authors:** Heather C. Etchevers

## Abstract

The goal of lineage tracing is to understand body formation over time by discovering which cells are the progeny of a specific, identified, ancestral progenitor. Subsidiary questions include unequivocal identification of what they have become, how many descendants develop, whether they live or die, and where they are located in the tissue or body at the end of the window examined. A classical approach in experimental embryology, lineage tracing continues to be used in developmental biology, stem cell and cancer research, wherever cellular potential and behavior need to be studied in multiple dimensions, of which one is time. Each technical approach has its advantages and drawbacks. This chapter, with some previously unpublished data, will concentrate non-exclusively on the use of interspecies chimeras to explore the origins of perivascular (or mural) cells, of which those adjacent to the vascular endothelium are termed pericytes for this purpose. These studies laid the groundwork for our understanding that pericytes derive from progenitor mesenchymal pools of multiple origins in the vertebrate embryo, some of which persist into adulthood. The results obtained through xenografting, like in the methodology described here, complement those obtained through genetic lineage tracing techniques within a given species.

## Introduction

### Brief history of the mysterious pericyte

The word “pericyte” carries its own uncertain definition - by its position, rather than by its function. Careful observations by Rouget and drawn using a camera lucida in 1873, showed highly ramified cells, which he considered likely to be contractile smooth muscle cells (SMC), apposed tightly to the hyaloid microvasculature of the frog and the rabbit [1]. Capillaries, apparently the simplest blood vessels, are not functionally identical around the body. What is in common is the unit of the endothelial cell, in direct contact with the circulating blood and in tight contact with its lateral neighbor. However, even endothelial cells can be differentiated molecularly and phenotypically based on their position in new sprouts, or in lymphatic beds. Most capillaries are associated with some form of pericyte, a multipotent contractile cell type on the immediate abluminal surface of the endothelial cells. These are responsible for the secretion of a tissue-specific basal lamina [2] and may be closely apposed or not [3]. Depending on vessel type, there may then be one or more concentric layers of SMC, all encased by outer connective cells including fibroblasts, which maintain the position of the blood vessel within the tissue or cavity.

The term pericyte has expanded over time to encompass the description of any resident periendothelial cell, though it tends to be applied most to microvessels. There are both molecular and phenotypic features that are either distinct to, or in common between, the following cell types: brain capillary pericytes, arterial, venous or lymphatic periendothelial SMC, kidney podocytes, coronary vascular pericytes, liver stellate cells, myofibroblasts, bone marrow stroma, dental pulp stem cells and even a subpopulation of macrophages [4, 5].

Many of the specific cell types listed above appear to retain mesenchymal multipotency into adulthood in multiple species, further confounding terminology. It is therefore essential to define the cell type under study with both molecular markers and in vivo position in tissue relative to vascular endothelium and, optimally, its basal lamina [6]; short of these conditions, it is important to keep in mind that all markers used to date are typical of, but never restricted to, a given pericytic population [7, 8]. For example, the large transmembrane glycoprotein encoded by the *CSPG4* (chondroitin sulfate proteoglycan 4) gene, also known as NG2 (for nerve/glia antigen 2), is expressed by many multipotent cell types, such as mesenchymal, keratinocyte and radial glia stem cells, in addition to pericytes.

### An adaptable tool: xenografting

A major fate-mapping technique to track cellular lineages [9], used in experimental embryology since its development in the last third of the 20th century, has been the construction of quail-chick chimeras. This approach exploits species differences in nuclear structure to permanently mark cells grafted from a donor to a host embryo [10], and has been adapted to highlight immunological differences with species-specific antibodies [11]. The flexible concept has taken many forms over the years. For example, mouse-chick and duck-chick chimeras have also been constructed, to study aspects of tooth or beak shape development and take advantage of introduced genetic modifications or species-specific attributes [12, 13]. Transfected quail donors have also made it possible to combine persistent fate-mapping with localized *in vivo* responses to changes in gene expression [14].

The endothelial cell lineage differentiates itself from other future mesodermal progeny at a very early time point, when the future head is barely distinguished by an anterior transverse buckling in the germ layers and gastrulation is still underway. The tyrosine kinase-linked receptor to the major bioactive form of vascular endothelium growth factor (VEGFA), Vegfr2, is already expressed then in a subset of cephalic mesoderm that subsequently differentiate into endothelial cells [15]. VEGF is essential to early vascular development and endothelial survival, and is secreted by cell, including pericytes, contacting sprouting neovessels under both normal and pathological conditions [16-18]. Xenografts of mesodermal mesenchyme lateral to the future brain at times before neural crest migration participate in both striated muscles and the endothelium of all cephalic blood vessels [19, 20]. Vegfr2-expressing precursors coalesce and cavitate to create a primary capillary network. Thereafter, new blood vessels sprout from pre-existing vessels. The growth process is known as angiogenesis and occurs throughout life and the body. In vertebrates, such labile sprouts are stabilized by pericytic contact and molecular cross-talk that appears to be evolutionarily conserved [16, 18, 21].

The development of antibodies against species-specific epitopes present on vascular endothelium, such as MB1/QH1 for the quail, which is not present in chicken endothelium [22, 23], prefigured the development of multiple conditional yet indelible fate-mapping techniques to trace the origins of endothelial cells from various intra-and extra-embryonic mesodermal sources. Confirmation of the mesodermal origins and further molecular properties of the vascular endothelium have been examined by fate-mapping in xenografting experiments [24, 25]. In the 2000s, Cre-lox [26] and Tol2 [27, 28] technologies allowed localized, timed recombination and subsequent expression of lineage tracers such as beta-galactosidase or fluorescent proteins, as well as functional manipulation of other genes’ expression in specific populations. The *Tie2* (*Tek*)-driven Cre recombinase has been particularly useful in following the vascular endothelial lineage in the mouse [29, 30] and validating the evolutionary conservation of the embryonic origins of the endothelium, but many other early and specific markers, such as *Vegfr2* promoter-driven fluorescent markers, are also used for lineage tracing in multiple model species like zebrafish [31].

As VEGF, BMP and related signaling pathways have been shown to be critical for the establishment of vascular identity and function in other parts of the body [29, 32], mediation by perivascular cells of signals between the surrounding tissue and the endothelium seems to be the rule and not the exception, no matter the embryological origin of the pericyte [33]. Because the potential for conferring these tissue-specific properties are at least in part a function of their origin, the results of pericytic lineage tracing remain relevant for understanding the causes of, and proposing adapted therapies for, vascular tumors and malformations and inappropriate angiogenesis.

### Multiple origins of pericytes - Neural crest

Neural crest cells (NCC) detach from the left and right boundaries between the ectoderm and the neural plate, as the latter rolls up into a tube which will give rise to the central nervous system. Fate maps using avian embryo chimeras have shown that these NCC migrate into and through the perineural mesodermal mesenchyme until they both mingle with it and colonize the appropriate distal targets, where they differentiate into the peripheral nervous system, certain types of endocrine cells, and all extra-retinal pigment cells (reviewed in Le Douarin et al. 2008). Cephalic NCC also engender many tissues that in the body are derived from the mesoderm, with the notable exception of vascular endothelium, derivatives grouped under the heading of the “mesectoderm”. These include the connective components of all head and neck glands, facial muscles and tendons. The dermis and adipose tissue overlying the jawed facial skeleton and brain case, the bones of that part of the skull; the meninges, including the vascular pia mater, underlying it are also of neural crest origin, as shown by lineage-tracing techniques in the avian embryo and later validated in other vertebrates [34-42].

Normally, NCC as well as Vegfr2-expressing mesenchyme surround the forebrain from the dorsal and ventral sides, respectively. Where they meet, they combine to form a leptomeningeal vascular plexus by the second day of incubation in the chick, although capillary penetration of the forebrain only occurs during the fifth day. In embryos experimentally deprived of rostral NCC, forebrain apoptosis occurs on the second and third days of incubation, such that the prospective forebrain tissue is not even present on the fifth day [43]. The experimental phenotype is due to a survival effect of mesenchyme of NCC origin on the early forebrain neuroepithelium within the pia mater, not compensated by unstabilized vascular endothelial precursors alone [44]. Much but not all of the NCC-derived perivascular mesenchyme in this location, both on the surface or intraparenchymally, ultimately expresses alpha-smooth muscle actin (aSMA). The migrating NCC normally counter environmental signals of the bone morphogenetic protein (BMP) and Wnt families and stimulate VEGF signaling [reviewed in [26]] by producing multiple members of the DAN (for differential screening-selected gene aberrative in neuroblastoma) family of antagonists: Gremlin (Grem), Dickkopf (Dkk1) and Cerberus (Cer1) [45, 46]. Combinations of both quail-chick chimeras and molecular biology have demonstrated that the anteroposterior level of NCC origin determines its potential to enable forebrain survival and growth [47], and that only NCC with mesectodermal potential normally differentiates into pericytes and SMC.

The first indications of the unique role of NCC in cephalic blood vessels came from fate-mapping experiments using radioactive isotopes, retroviral infection or xenografting to show their constitution of the branchial arch mesenchyme and subsequent incorporation into the muscular walls of the corresponding large arteries [35, 48, 49]. NCC derived from the posterior rhombencephalon were shown to contribute all components of the proximal large arteries to the heart, with the exception of the endothelium. Subsequently, the functional role of NCC of this origin was noted in the septation of the pulmonary trunk from the aorta [50, 51]. NCC are equally important to the smooth muscle wall of the posterior aortic arch arteries in mammals [26] and integrate into the walls of the cardinal veins [49, 52]. NCC were shown with quail-chick chimeras to provide two distinct types of cells in the tunica media of large elastic arteries derived from the aortic arches, some non-SMC adventia, and an early periendothelial layer of NCC not organized in lamellar layers, and only sometimes expressing aSMA [53].

#### Abutting pericyte lineage-tracing

The cellular resolution afforded by interspecies chimeras enabled discovery of the origin and extent of a subpopulation of pericytes in the multipotent cephalic NCC. Isotopic xenografts of small fragments of neural folds from brain levels corresponding to the future diencephalon and mesencephalon of quail donors resulted in abundant graft-derived NCC within the forebrain meninges of the chicken hosts, with a sharp border at the forebrain-midbrain boundary [52]. Quail cells, adjacent to endothelial capillary walls within the parenchyma and the leptomeninges, were co-immunostained with aSMA after a few days’ maturation and determined to be pericytes (Figure 1). The vascular media of veins and sinuses were only a grafted cell or two thick, confirming independent work also using quail-chick chimeras [49]. SMC and outer mural connective tissue cells of graft origin were organized in cell-dense layers in the distal portions of the major cephalic arteries; most of these in cross-section were concentric and interspersed with eosinophilic elastin. Histological differences in elastin organization between the parts of the aorta surrounded by SM of NCC *versus* mesodermal origin have also been noted by others [54], a heterogeneity much later borne out by functional studies of *ductus arteriosus* closure at birth in mice, as described below [55].

**Figure 1:**
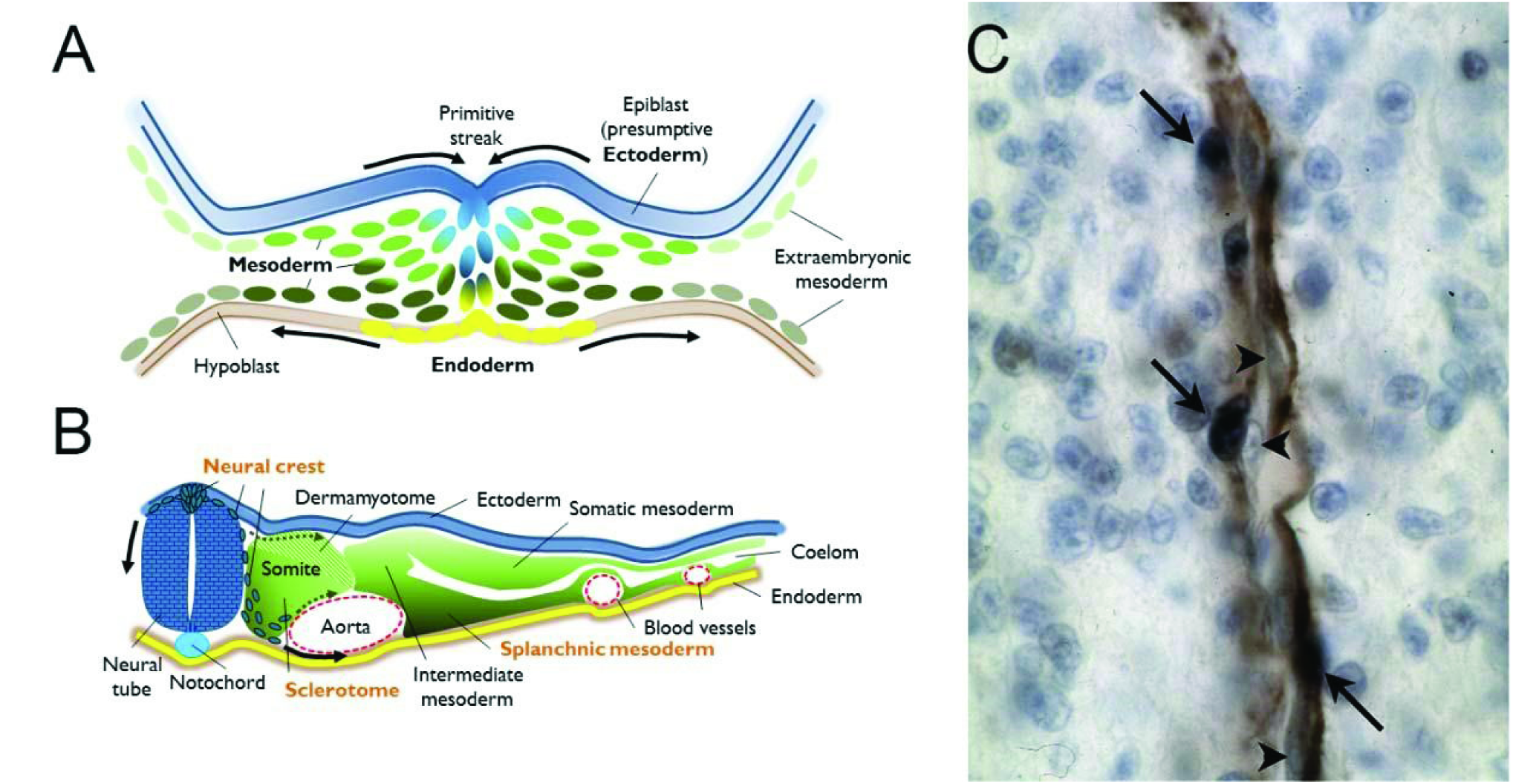
**A.** Drawing of a cross-section through the primitive streak of a vertebrate gastrula, highlighting contiguous populations of the three definitive germ layers (ectoderm, mesoderm and endoderm). These will each give rise to subpopulations with distinct potentials for cellular progeny by the neurula stage. **B**. Drawing of a cross-section through an idealized vertebrate neurula. The neuroepithelium has invaginated to detach from the ectoderm, in blue, and the neural crest cell population has been specified at their interface and begin to undergo an epithelio-mesenchymal transition (EMT) to migrate initially between the somite and the neural tube. This first ingression of cells is followed by streams migrating through mesodermal cell compartments, particularly the ventral somite as it also undergoes EMT to become the sclerotome. Both neural crest and sclerotome are sources of pericytes. Finally, the splanchnic mesoderm, the ventral compartment of the lateral mesoderm, is the source of both epicardial and other mesothelial cells which will provide the pericytes of heart and internal organs derived from the endoderm. **C**. Example of a homotopic graft of quail neural crest into a chicken host. Immunohistochemistry against a quail antigen (dark blue nuclei, arrows) highlights the presence of quail-derived pericytes among the alpha smooth muscle actin-expressing cells (brown) of a forebrain capillary. Arrowheads indicate unlabeled endothelial cells in close apposition. Photograph courtesy of the author.

Grafts of neural folds of the anterior rhombencephalon gave rise to all but the endothelial cells of the proximal segments of the arteries irrigating face and forebrain, as well as the distal internal carotid arteries. Median rhombencephalic neural folds contributed cells to the carotid arteries and cardinal veins, while posterior rhombencephalic NCC tended to incorporate into proximal segments of the carotid arteries, the aortic and pulmonary trunks and the conotruncus and semilunar valves of the heart itself, confirming reports by others [52, 56]. Thus, the original anteroposterior origin of a cell within the neural folds corresponds to its final distoproximal distribution in a defined subset of cephalic blood vessels. This subset, designated the “branchial vascular sector”, is a distinct circuit of blood vessels originating in the ventral aorta and aortic arches, ramifying into circumscribed capillary plexuses, and terminating in their venous return to the heart. These vessels irrigate the forebrain, neck, face and jaws, and in what appears to be a species-dependent pattern, anastomose with the evolutionarily more ancient vascular network in the head [52].

Pdfgrb appears to play an evolutionarily conserved role in pericyte biology in microvessels, since its promoter driving a fluorescent marker has enabled lineage tracing and live imaging of vascular mural cells within an elegant zebrafish model [57]. In order to observe the dynamics of pericyte-EC interactions, EC can be simultaneously traced with a distinct fluorescent reporter [58], or *in vivo* after lipophilic, fluorescent dye injection into chicken host veins [59]. It is possible to make chimeras by transplanting cells from labeled donors to unlabeled wild-type hosts, for example with transgenic GFP-expressing chick embryos, or with transgenic zebrafish particularly between blastula and gastrula or shield stages, where detailed fate maps are available. Chimeras can thus be a vital tool for dissecting questions related to cell autonomous or non-autonomous effects of endogenous or exogenous signaling molecules on specific cell lineages, in combination with genetic gain-or-loss of function. To date, however, this labor-intensive strategy has been eschewed in favor of genetic combinations of lineage-specific labels with respect to studying teleost pericytes. This has been fruitful in confirming that the evolutionarily more recent brain and face indeed have neural crest-derived pericytes, while the rest of the body has mesoderm-derived pericytes, as shown by the reporters driven by the transcription factor genes *Sox10* and *Tbx6*, respectively [57].

### Multiple origins of pericytes - Mesoderm

A second vascular division can be distinguished in the head by virtue of its vessels *not* belonging to the branchial sector. Like in the rest of the body, its component arteries and veins share the property of being entirely constructed from the embryonic mesoderm. The cephalic mesoderm, although mostly mesenchymal, is analogous to the axial and paraxial mesoderm of the body. In mature vertebrates, vessels branching rostrally from the aorta irrigate the dorsal head, including the midbrain, cerebellum and hindbrain. This vascular domain contacts the branchial sector at the circle of Willis, a large anastomosis between the bifurcation of the basilar artery and the cerebral arteries, branches of the internal carotids. This polygon surrounds the optic nerves and ventral diencephalon, reflecting the transition (at least in birds) within the meninges from an entirely mesoderm-derived region, the midbrain, to a composite mesoderm/NCC-derived region, the forebrain [52].

#### Epicardium

At the level of the heart, there is another vascular interface of neural crest-derived pericytes and SMC with their mesoderm-derived counterparts. The epicardium is a thin layer of specialized mesothelium that spreads from cells clustered on the coelomic surface of the right *sinus venosus* over the dorsal wall of the heart tube and toward the outflow tract during cardiac morphogenesis. A replication-defective adenovirus carrying the *LacZ* gene driven by a strong promoter, was used to follow the fate of a subset of either chicken proepicardium directly infected *in ovo* [60] or cultured quail epicardial cells, or whole quail proepicardial primordia grafted into chick hosts [56, 61]. In all cases, cells underwent an epithelial-mesenchymal transition in order to leave the epicardial layer and integrate into the tunica media of both coronary arteries and veins. In addition, this epicardially derived mesenchyme also generated interstitial fibroblasts amongst the myocardium, indicating that this alternate source of pericytes and smooth muscle retains the capacity to give rise to non-vascular connective tissue, like its neural crest counterpart in the head. Some of the endothelial cells of the coronary vasculature were also graft-derived in independent studies [56, 60], indicating that either it is difficult to entirely remove endothelial components from grafted tissues (a potential pitfall for chimera approaches), or that within the mesoderm, contrary to the neural crest, endothelial cells and vascular pericytes share a common lineage. Certainly, pericytes and coronary artery SMC do, and fate-mapping in mice demonstrated that the first is a potential resident progenitor pool for the second, when it is possible to remobilize them pharmacologically [62]. Recently, at least some coronary arterial pericytes and smooth muscle have also been fate-mapped to the hindbrain neural crest in both mouse and chick [63], implying that the heart is also a zone of transition from NCC-derived to mesoderm-derived pericytes.

#### Mesothelium

The part of the lateral plate mesoderm that is closest to the endoderm and its signals is the splanchnic mesoderm. Within this original two-dimensional epithelium, flanking the axis, are fields corresponding to distinct populations of prospective mesenchyme. In gastrulae, the anteriormost zone of embryonic “lateral” mesoderm beyond the future head will integrate into the liver, followed by the inverted U-shaped first heart field overlying the anterior intestinal portal [64], which will give rise to the left ventricle. This zone is succeeded in posterior order by the splanchnic mesoderm of the future second heart field (SHF), which follows the same shape slightly more median. At the posterior ends of the U, beyond the cells that will integrate into arterial pole myocardium [65], only the right primordium is maintained from initially bilateral prospective proepicardial fields, at least in amniotes. Chicken chimeras hosting grafts of vital dye-labelled lateral plate mesoderm subpopulations demonstrated that medial somatic mesoderm (in contact with the ectoderm, dorsal to the right sinus venosus) also contributes to the proepicardium [66]. All of these zones are capable of generating vascular SMC and fibroblasts, while only the posterior SHF [67] and epicardium make unambiguous pericytic contributions.

Further posterior, the splanchnic mesoderm gives rise to the *Wt1*-expressing mesothelial layer covering all coelomic visceral organs – particularly the liver and digestive tract, but also the kidneys and gonads. This mesothelium contributes mesenchymal cells expressing aSMA and desmin, clearly SMC within arteries and veins but on smaller microvessels also, to the invasive vasculature of the entire developing gut [68].

#### Sclerotome

In counterpart, grafts of the presomitic mesoderm from quail to chick hosts have demonstrated that not only endothelial cells but also all the pericytes and vascular SMC of the limbs and trunk are derived from this paraxial population [69]. The lack of contribution to vascular SMC of the viscera was also noted in this work, indirectly confirming its origin from the splanchnic mesoderm. Furthermore, it is the part of the paraxial mesoderm in contact with the endoderm, which after first epithelialization into the somites, later becomes mesenchymal, that is the source of these outer body wall pericytes and vascular SMC. The pericytes were embedded in a laminin-rich basal lamina, and within the aortic wall, came from the Foxc2-expressing sclerotome. Interestingly, while Pdgfrb-positive grafted cells were numerous within the aortic SMC, the periendothelial cells expressing aSMA most strongly did not express as much (or any) Pdgfrb as the more peripheral layers of the tunica media [69]. Further work showed that these sclerotomal smooth muscle cells had been preceded by a temporary contribution from the splanchnic mesoderm [70].

One might therefore predict that quail-chick chimeras of the ventral intermediate mesoderm, between paraxial and lateral plate populations, may also demonstrate pericytic and smooth muscle potential, perhaps for the specialized kidney pericytes known as podocytes that can undergo pathological fibrosis in response to injury [71]. Extraembryonic mesoderm, beyond but still continuous with the lateral plate, also may have pericytogenic potential for the vascularized extraembryonic membranes [72].

Hereafter, I will describe steps to track neural crest-derived pericytes in and around the forebrain. The protocol is highly adaptable and has been often described for whole neural tube grafts, but can be applied to any embryonic or extraembryonic tissue that can be physically manipulated.

## Materials

### 2.1 Eggs

Freshly-laid fertilized chicken and quail eggs that have been stored for under a week at 12-15°C (approximately 55-60°F) are critical for the chimeric embryo technique. Viability and normal development rapidly decreases in the population thereafter, although individual eggs may still develop well up to 10-14 days of cool storage.

### 2.2 Incubators and handling

1. A standard wine cooler-type refrigerator can be adapted to maintain the appropriate temperature for cool storage, as can a ventilated laboratory incubator that incorporates both refrigeration and heating elements to maintain a constant temperature.
2. The warm egg incubator should have forced air circulation as opposed to passive temperature control, to ensure that all parts of the interior have the same temperature and humidity. It should also be equipped with a hygrometer. Once development has started by warming the eggs, they must be maintained at 38 +/- 1°C (98-102°F) and 50-60% humidity. It is preferable to err on the cooler rather than the warmer side to reduce risk of increased mortality and malformations.
3. During cool storage, eggs may be kept upright. Replaceable cardboard or styrofoam egg crate trays in which eggs are delivered are adequate to incubate eggs on the side 24h before opening. Plastic, plexiglass or varnished wooden trays approximately 1 cm (1/2 inch) thick, 5 cm (3 inches) wide, and as long as the incubator, with oval holes about 3 cm long (a little more than an inch) to secure eggs on their sides in the depressions, are useful. These allow the researcher to supervise development of dozens or hundreds of eggs after sliding trays out every couple of days, to remove any that have contracted an infection or have died. Single segments of such trays make useful holders for securing individual eggs during the microsurgical procedures and are recyclable.
4. To make cephalic neural fold grafts at stages HH8 [73], optimally at 3-6 visible somites, place chicken eggs for approximately 27-30 hours and quail eggs for 24 hours in the 38°C incubator.

### 2.3 Microdissection tools

Microsurgery tools should be regularly sterilized in a dry oven for 2 h at 150°C with appropriately resistant silicon protection, or 20 minutes at 120°C in an autoclave. Allow to cool before use.

Useful tools include

- One pair of sharp-tipped, 4-inch, curved microdissection scissors
- One pair of blunt-ended dissecting forceps, 0.8 mm tip width (coarse)
- One pair of Dumont no. 4 dissecting forceps, 11 cm long, 0.13 X 0.08 mm tips (medium)
- Two pairs of Dumont no. 5 dissecting forceps, 11 cm long, 0.1 x 0.01 mm tips (“biologie”)
- One perforated stainless steel spoon for gentle embryo transfer
- One pair of small Vannas or Pascheff-Wolff ophthalmological spring scissors
- Two screw-chuck dissection needle holders with stainless steel Minutien pins (diam. 0.2 mm, 1.2 cm length), pre-sharpened in oil on a fine-grit Arkansas stone and cleaned, or electrolytically sharpened tungsten knives [74]
- Optional substitute: a Dean microdissecting knife; curved; 1 x 7 mm blade
- Prepare borosilicate glass micropipettes on a conventional electrode puller or from borosilicate Pasteur pipettes over a flame, in order to generate micropipettes with tapered tips of about 2-3 cm long. Break the tips, using appropriate eye and hand protection from possible fragments, by bending with forceps to obtain a round orifice with a diameter of approximately 100-200 microns. These may be prepared in advance if sheltered from dust and humidity.

### 2.4 Other equipment

- Aspirator tube assembly consisting of an inflexible plastic mouthpiece such as a truncated sterile pipette cone with filter; at least 40 cm of latex tubing (5 mm internal diameter, 7 mm outer diameter); and if using micropipettes, a flexible silicone rubber nosepiece
- Stereomicroscope (e.g., Leica MZ 7.5)
- Swan-neck fiber optic illumination
- Sterile glass 50-60 mm dissection dish lined with several millimeters of silicone base such as Sylgard^®^ (*see* **Note 1**)
- Sterile tissue culture dishes (glass or plastic), 35 mm, or watchglasses
- Syringes: 1 mL, 2 mL
- Needles: ½ inch, 26 gauge; 1 inch, 18 gauge.
- Plastic or flame-polished standard Pasteur pipette and bulb
- 5 cm (2 inch) wide thin, transparent packing tape (*see* **Note 2**).
- Glass Coplin jars

### 2.5 Solutions

- Phosphate-buffered saline (PBS), without additional Ca^2+^/Mg^2+^
- Penicillin-streptomycin 100X for tissue culture - 0.5 mL aliquots kept at −20°C.
- Prepare the following just before beginning microsurgeries: 50 mL PBS in a sterile tube with 1X final concentration penicillin-streptomycin (AS-PBS).
- Dilute one part carbon-based, opaque India drawing ink (also called Chinese ink) to 4 parts AS-PBS (*see* **Note 3**).
- Modified Carnoy’s solution:

- 11% formaldehyde (3 parts commercially available 37% solution)
- 10% glacial acetic acid (1 part)
- 60% ethanol (6 parts)
- 70%, 90% and anhydrous 100% ethanol
- Toluene or xylene (*see* **Note 4**)
- Permanent, solvent-based resinous mounting medium
- Antibodies: QCPN (*see* **Note 5**) or QH1 supernatants (*see* **Note 6**); widely commercially available ascites from the 1A4 clone of mouse monoclonal antibody to aSMA (*see* **Note 7**); anti-mouse-IgG1 conjugated to horseradish peroxidase (anti-IgG1-HRP); anti-mouse-IgG2a conjugated to alkaline phosphatase (anti-IgG2a-AP).
- 50 mM or 0.15% w/v glycine in 50 mM ammonium chloride in distilled water
- PBT: PBS with 0.1% Tween-20 detergent from a 10% stock kept at 4°C
- Heat-inactivated goat or fetal bovine serum
- 10% or 30% H_2_O_2_ (stock, store in dark at 4°C)
- TBST: for 1L total volume with distilled water, use 25 mL 1M Tris-HCl, pH 7.5; 20 mL 5M NaCl; 0.2g KCl; and 10 mL 10% Tween-20.
- NTMT: for 500 mL total volume with distilled water made extemporaneously, use 25 mL 1M Tris-HCl, pH 9.5; 10 mL 5M NaCl; 25 mL 1M MgCl_2_; 5 mL 10% Tween-20; 0.1g levamisole.
- 0.5 mg/mL 3,3-diaminobenzidine tetrahydrochloride (DAB), prepared in PBT just before use from frozen aliquots of 10 mg/mL stock solution in PBS (*see* **Note 8**)
- 75 mg/ml nitroblue tetrazolium (NBT) in 70% dimethylformamide
- 50 mg/ml 5 bromo-4-chloro-3-indolyl phosphate (BCIP) in dimethylformamide.

## 3. Methods

### 3.1 Host egg preparation

Incubate host embryo horizontally overnight and mark the highest point on the side of the egg with a short pencil mark (*see* **Note 9**). Scrub down the top surface of each egg briefly with a paper towel moistened in 70% ethanol; do not use soap and water or wet the shell directly, as it is porous and viability can be affected. In order to allow the embryonic disk to disengage from the upper eggshell, gently drill a small hole in the large end of the egg with the curved scissor tips. Insert 18-gauge large-bore needle attached to a 2 mL syringe, oriented toward the bottom of the egg, and remove approximately 2 mL of liquid albumin. The albumin close to the yolk is thicker and if it blocks the needle, expel again and orient the point elsewhere in the egg; if any yolk enters the needle or syringe, it has been pierced and the egg should be discarded. Tape the hole with a small piece of transparent tape. Gasses coming through the eggshell will equilibrate the lower internal pressure with the atmosphere and enable the yolk to detach from the shell surface.

With gentle pressure, next insert the lower point of the scissors and cut a small circular window into the air pocket of approximately 2-2.5 cm diameter around the pencil mark. Smaller gestures generate more fragments but better control of the curve of the window. Pry the circle up at the end of the cut to remove without allowing surface shell to fall into the egg. Carefully pick out any small pieces that have fallen within using blunt forceps, and rinse with approximately 0.5 mL antibiotics-supplemented PBS, using the large blunt-ended Pasteur pipette, withdrawing most of the liquid and replacing with a few drops of clean AS-PBS.

Fit a ½ inch, 26 gauge needle to a 1 mL syringe. Press the needle within the loosened cap to introduce an angle of about 30-45°, with the orifice facing the inside of the angle. Fill the syringe with dilute India ink (*see* **Note 3**). Use needle to pierce yolk at edge of blastoderm, orienting upwards towards ventral side of host embryo. Gently inject approximately 0.2 mL without introducing air bubbles, and shake horizontally to spread over yolk beneath the blastoderm, providing contrast.

Using Dumont no. 4 dissecting forceps, pinch the vitelline membrane lateral to the head fold, without catching the underlying embryo. Raise slightly and move to the side to pull a hole in the membrane over the head, releasing at a more caudal level. Gently add a couple of drops of AS-PBS into the hole to help separate the vitelline membrane from the underlying embryo.

### 3.2 Host microdissection

Neural folds between the levels of the presumptive diencephalon and medulla, facing the fifth somite pair, produce neural crest cells that later give rise to pericytes and vascular smooth muscle in predictable tissues of the head, neck and heart [35, 49, 52, 63, 75-77]. The forebrain, midbrain and first rhombomere lend themselves best to neural fold grafts at 3-5 somite pairs, before closure, rather than the entire tube. Depending on the desired experiment, unilateral or bilateral grafts may be made at these levels.

With a microscalpel or blade, make horizontal cuts into the neural folds at the rostral and caudal limits of the area to be removed and grafted. At this stage the folds spanning the transition from outer ectoderm to inner neuroectoderme are raised above the mesoderm; superficial “tickling” motions, with quick upward strokes of the point of the blade, are effective to cut through the single cell layer without penetrating too deeply. Lateral cuts on first the ectodermal, then the neuroectodermal faces of the epithelial folds will permit a rectangle of tissue to be nudged rostrally and off the underlying mesoderm. If the cuts attain the mesoderm, the drag on this mesenchyme may perturb the underlying endoderm, which can induce either collateral malformations or death.

A gentle drop of AS-PBS over the embryo, not directly on the area to be grafted, and either a Petri dish lid or a watchglass, will protect the host until grafted tissue can be placed into it. For more than fifteen minutes’ wait, replace the embryo in the incubator.

### 3.3 Graft egg preparation

Crack a GFP-chick or quail donor egg into a 10 cm or a 6 cm Petri dish, respectively. Use a corner of a piece of paper towel folded into a 2.5 cm (1 inch) square to wipe the thick albumin radially away from the light blastodermal disk. After adding a couple of drops of AS-PBS to the surface, use the microdissection scissors to cut into the yolk around the blastoderm and the central embryo. Catch a corner of the circle with the medium forceps and bring the perforated spoon superficially beneath, attempting to slide the embryo onto the spoon while bringing as little yolk as possible. Transfer into a 60 cm Petri dish filled with AS-PBS and jiggle to remove the milky yolk on the ventral side and the vitelline membrane. Use the perforated spoon to transfer into the dissection dish with clean AS-PBS. Immobilize by using the medium forceps to pin out the corners with Minutien pins.

### 3.4 Graft microdissection

For rostral neural folds at 4-5 somite pairs, prepare the grafts by cutting out the equivalent areas as described above in 3.2 to transfer to the host. For later or more caudal grafts of whole neural tube, or somites, it may be necessary to isolate the tissue and remove any potentially adherent mesenchyme with pancreatin [59, 78] (*see* **Note 10**). Distinguish the rostral from caudal end by notching a corner or introducing a slightly trapezoidal shape.

Using the mouth micropipette for fine control of the piece, first rinse by aspirating and expelling a little yolky AS-PBS. This will prevent the graft from sticking in the pipette. Aspirate the graft and some microliters of AS-PBS by orienting the micropipette toward the thin end of the rectangle in the dissection dish. Avoid scraping the tissue against the sharp edges of the pipette (*see* **Note 11**).

Transfer the graft slightly lateral to its final placement so that the liquid drains off, and use the microscalpel to gently nudge it into the same orientation as it came from originally, before sliding into position. Aspirate excess AS-PBS from near (but not over) the graft into the micropipette. This will bring graft and host tissues into close contact under the meniscus and aid in healing.

Seal the window with tape, ensuring that no folds remain along the edge that would allow direct air entry to the egg (or albumin seepage out). Replace horizontally in warm, humid incubator for desired interval. For cerebral pericytes, initial investment of the forebrain meninges is visible from two days later and ingression of grafted aSMA-positive cells into the parenchyme, three days later and beyond.

### 3.5 Harvesting, fixation and processing for embedding

1. Make a slit in the yolk above and below the live embryo to be removed (*see* **Note 12**). Use a Pasteur pipette to drip some PBS below, to help detach it from yolk.
2. Grasp a corner of the *area opaca* with medium forceps and continue to hold while snipping the lateral cuts to detach embryo entirely.
3. Still holding the corner of the *area opaca*, bring slotted spoon underneath to transfer embryo to a dish with clean PBS.
4. Remove all traces of ink, yolk, and extraembryonic membranes.
5. If helpful, pin out embryo or desired organs before fixing on a dedicated dissection dish, not to be also used for graft preparation. Fix for 1h to overnight in modified Carnoy’s solution, the timing as an empirical function of size (see **Note 13**).
6. Rinse abundantly in 70% ethanol (EtOH) (see **Note 14**).
7. Dehydrate in 90% then at least two changes of anhydrous 100% EtOH. Keep recipients closed, as the 100% EtOH can be hygroscopic, to the detriment of section quality later in the process.
8. Clear thoroughly in toluene, xylene or a limonene-based clearing agent (see **Note 4**).
9. Substitute with three changes of melted Paraplast^®^ X-tra in a heat-resistant recipient in an oven at 55°C for a period of hours (HH10) to overnight (after HH17), to be determined empirically.

### 3.6 Immunolabeling

It is important to choose combinations for co-labeling that allow for isotype discrimination and optimal contrast in labelled components. For example, the monoclonal antibodies QCPN (all quail calls) and QH1 (quail endothelial cells) are mouse isotype IgG1, while aSMA is mouse isotype IgG2a (*see* **Note 15**). In this example, we perform immunohistochemistry against quail cells with QCPN and against aSMA to highlight differentiating perivascular cells among them, in sections from rostral head after the grafts performed above at HH8 and harvested at HH25 [73]. All steps are performed at room temperature unless otherwise indicated.

1. Deparaffinate 5μm-thick microtome sections in Coplin jars with three changes of five minutes each of xylene (*see* **Note 4**).
2. Rehydrate with two changes of two minutes each of 100% anhydrous EtOH, followed by two changes of two minutes each of 90% EtOH, and one immersion for two minutes each in 70%, 50%, 30% EtOH before bringing slides to distilled water.
3. Incubate 10′ in freshly prepared glycine/ammonium chloride solution (*see* **Note 16**).
4. Rinse twice in PBT then place in PBT with 3% final volume H_2_O_2_ for 10 minutes.
5. Rinse three times in PBT, then further block non-specific protein-protein binding with 2% goat or bovine serum in PBT for 30’ under coverslips (*see* **Note 17**).
6. Gently remove coverslips, blot excess liquid, and replace with primary antibody mix. Here, we foresee 100 μL per slide and make up a dilution of ½ QCPN supernatant (*see* **Note 18**), 1/400 aSMA, 1/50 goat or bovine serum and the rest of the volume with PBT. Apply coverslips to slides and place in humid chamber. Incubate 2h or overnight at 4°C in a humidified chamber (*see* **Note 19**).
7. Remove coverslips, blot excess liquid, then wash slides in Coplin jars filled with PBT 5 times for at least 5 minutes.
8. Make up a dilution of 1/400 anti-IgG1-HRP, 1/50 goat or bovine serum and the rest of the volume with PBT. Apply 100 μL per slide under coverslips, and place horizontally in humid chamber for 1h.
9. Remove coverslips, blot excess liquid, then wash slides in Coplin jars filled with PBT 3 times for at least 5 minutes.
10. Dilute DAB in PBT and add 0.003% H_2_O_2_, before immerging slides and keeping in dark for approximately ten minutes. Wear gloves. Rinse in PBT before examining signal intensity, which should show dark brown grafted nuclei on a white background. Stop reaction by rinsing all slides twice more in PBT and once in TBST.
11. Make up a dilution of 1/400 anti-IgG2a-AP, 1/50 goat or bovine serum and the rest of the volume with TBST. Apply 100 μL per slide under coverslips, and place horizontally in humid chamber for 1h.
12. Remove coverslips, blot excess liquid, then wash slides in Coplin jars filled with TBST 5 times for at least 5 minutes.
13. Wash slide in freshly made NTMT twice for at least 5 minutes. Alkaline pH is very important at this step.
14. Using 0.5 μL NBT and 3.5 μL BCIP stock solutions per mL NTMT, immerge slides and keep in dark. Rinse in NTMT to examine signal every 10 minutes until arteries begin to develop a bluish-purple circumference. Stop reaction by rinsing excess NBT-BCIP solution and then immerging in PBT.
15. Rinse in PBT and then apply glass coverslips over AquaTex ^®^ mountant.

## Notes

1. Opaque black-tinted silicone is preferable to transparent, for better contrast.
2. The transparency is only useful for peeping through the egg window within the confines of the incubator to see if the embryo is well vascularized and, at later stages, showing spontaneous movements. At room temperature, the window rapidly fogs over with condensate, so opaque tape can work also.
3. Pre-test the ink preparations alone on eggs, as different brands may be prepared in differently toxic diluents. Rare brands of packing tape also emit noxious odors that can be a source of reduced viability.
4. The advantages of various clearing agents, be they solvants, terpenes or other substances, are a subject of great debate [79]. We have successfully used toluene, xylene and limonene-based Histoclear^®^ to clear tissues as well as to permanently mount sections for light microscopy.
5. QCPN was deposited to the Developmental Studies Hybridoma Bank (DSHB) at the University of Iowa, U.S.A., by J. A. and B. M. Carlson (http://dshb.biology.uiowa.edu/quail-cell-marker).
6. QH1 was deposited to the Developmental Studies Hybridoma Bank at the University of Iowa, U.S.A., by F. Dieterlen-Lièvre (http://dshb.biology.uiowa.edu/endothelial_2).
7. Alpha-smooth muscle actin monoclonal antibody [80] is now widely distributed by commercial suppliers.
8. DAB, NBT, BCIP and dimethylformamide are all biohazardous substances requiring the use of gloves and eye protection.
9. All eggshell labeling should be made in pencil as opposed to ink, to avoid potentially toxic solvent exposure.
10. Videos of two applications of the avian chimera technique, described in an earlier protocol relative to whole neural tube grafts [78], are freely available at https://www.jove.com/video/52514/ and https://www.jove.com/video/51534 [59, 81].
11. For larger pipette diameters, holding the tip a second in the flame after breakage can fire-polish (round) the potentially damaging edges.
12. Dead or dying embryos rapidly undergo necrosis and are never worth processing. Viability is easily visible after HH15 in the vibrant red of the blood in the vitelline veins around the embryo and the visible heartbeat.
13. 4% neutral buffered paraformaldehyde in PBS is a good generic fixative appropriate for either cryostat or paraffin sections; a minority of antibodies will not work after the solvents and heating denaturation inherent in paraffin embedding or antigen retrieval. The modified Carnoy’s solution used here preserves tissues more firmly than paraformaldehyde and is compatible with available QCPN, QH1 and aSMA antibodies. Tissues should be uniformly firm and white after fixation with modified Carnoy’s solution but will remain more flexible in paraformaldehyde. Overfixation by the former can lead to brittle, easily damaged embryos. Manipulate both fixatives under a fume hood with a charcoal filter to avoid damaging airway passages and the cornea. Rinse embryos after modified Carnoy’s fixative with 70% EtOH and paraformaldehyde with PBS.
14. Hospital pathology departments often have automated systems for tissue infiltration from ethanols to wax. These are fine, if the size of the tissue to be embedded is taken into account in the program and cassette chosen.
15. Directly conjugated antibodies can allow the use of other markers as well; we have successfully used FITC-conjugated aSMA and anti-fluorescein secondary antibodies coupled to enzymes or Alexa Fluor^®^-488 to convert to other chromogens or to reinforce resistance to bleaching in fluorescence microscopy. Immunohistochemistry is best used for two or more markers where the chromogens are chosen to have maximal contrast and are in distinct cellular compartments, yielding slides that are stable for years and require only a good light microscope for examination. Otherwise, immunofluorescence with optical sectioning or confocal microscopy is technically preferable in many ways. Durability is improved by the use of Alexa Fluor ^®^ fluorochromes with distinct emission spectra and a hardening mounting medium designed to preserve fluorescence, but costly imaging stations are required to acquire and record the results.
16. This step blocks free aldehyde groups from non-specific binding of antibodies and can reduce background signal as well as autofluorescence.
17. We cut hydrophobic “coverslips” for blocking and antibody incubations from rectangles of Parafilm M. Leaving the paper backing, foresee two coverslips per square outline. Cutting strips of (eg.) four squares for eight coverslips, and taking care not to crumple the film, peel back the paper longitudinally but only halfway across the strip, and fold back. It is then possible to cut across the strip at each horizontal line and again below the words “Laboratory Film”, leaving a paper “handle” on each coverslip to aid in removal of the backing. Remove paper entirely before positioning with coarse forceps over 100-200 μL of liquid on a 25x60mm slide surface.
18. Hybridoma supernatants have variable amounts of antibody present, depending on the production lot. We have also successfully used 1/50 dilutions of QCPN as provided by the DSHB.
19. This is a plastic box for storing 50 slides upright, with a waterlogged paper towel in the bottom and the slides balanced horizontally across the width of the column. It is placed in a plastic bag for overnight incubations.
20. Inactivate remaining DAB solution by oxidizing with dilute bleach before disposal, then rinse glassware well before reuse.

## Discussion

### Fate-mapping can further understanding of the pathophysiology of vascular malformations

Moya-moya disease (MMD) associates bilateral stenosis of the internal carotid or anterior/middle cerebral arteries, often at the level of the circle of Willis, and a profusion of telangiectatic blood vessels in a stereotyped distribution, allowing some bypassing of the stenotic area. The fact that MMD is restricted to this particular segment of the cephalic arteries, and that it has been associated with neurocristopathies, led to the hypothesis that a somatic mutation may manifest in the NCC pericytic component derived from descendents of a cell fate-mapped to the mid-rhombencephalic neural folds [82]. However, although the phenotype is localized, susceptibility can be dominantly inherited, although at low penetrance. Eastern Asian MMD is associated with a founder variation in the *RNF213* gene, encoding a ring-finger AAA-type ATPase with E3 ubiquitin ligase activity [83]. A wide variety of mutations in this gene are not only associated with MMD in other ethnic populations but with intracranial aneurysm in some [84]. Unlike the ectopic but localized angiogenesis of MMD, intracranial aneuryms show thinned SMC and elastic lamina within the arterial walls at the sites of dilation, without auxiliary vessel growth. Although its role in preventing aberrant angiogenesis, particularly in the head, appears certain from knockdown studies in zebrafish [83], it is unclear how the ubiquitous expression of *RNF213* translates into such a site-specific phenotype, or in which cell lineage (endothelial, NCC-pericytic, mesodermal-pericytic, or all of the above). Fate-mapping of each lineage, particularly using conditionally mutated animal models, could address these questions.

### Novel cellular contributions to secretory function

Pericytes of the microvasculature feeding the head and neck endocrine (pituitary, pineal, thyroid, parathyroid and ultimobranchial, for lower vertebrates) and exocrine (salivary, sweat, sebaceous, lachrymal) glands are also derived from NCC. While they are identifiably perivascular by position and alpha-SMA expression in the posterior pituitary gland, additional mesenchymal, non-endocrine NCC derivatives are present around and throughout the anterior hypophysis [35, 43, 52]. All of these cells, in the rat, appear to be nestin-immunoreactive [85], while a subpopulation of the anterior pituitary also expresses fibrillary collagen mRNAs and desmin, non-exclusive markers of pericytes [86]. Experimental removal of the neural folds leads to a malformed pituitary gland in chick [43]. It is unclear if the NCC act on the developing pituitary gland through their role as pericytes in stabilizing the meningeal and glandular capillaries, or if there is a more direct trophic effect on, or even cellular contribution to, neuroendocrine targets [87]. Like the retinal primordia, salivary and lachrymal glands, and the entire telencephalon, both Rathke’s pouch and the infundibulum of the diencephalic floor form initially, but are not maintained and undergo apoptosis in the absence of sufficient NCC [43, 47].

Thymic development is likewise aborted when NCC are experimentally removed [88]. In conjunction with malformations of the cardiac outflow tract, the effects of NCC ablation correlate with their normal differentiation into thymic vascular pericytes [35, 89], later mediators of CD4+ T-cell emigration into the bloodstream [90]. The association of effects on thymus and cardiac outflow tract development implicates NCC indirectly in the pathophysiology of DiGeorge syndrome [91].

### Pericytic mimicry and physiological stem cell niches

We and others have observed the normal presence of pigmented melanocytes in a pericytic position surrounding capillaries in the brain meninges and the pituitary of both rats and mice [92, 93], and in the *tunica media* of NCC-supported blood vessels immediately afferent and efferent to the heart [55]. In the latter situation, melanocyte precursors carrying a stabilized, overactive form of a downstream effector of Wnt signaling were fate-mapped in transgenic mice. Bipotent cells that could differentiate into either melanocytes or SMC, even though a minority among the SMC population of the embryonic arterial shunt called the *ductus arteriosus* (DA), leaned toward more melanocyte differentiation with Wnt signaling. By so doing, they thereby deprived this vital blood vessel of sufficient SMC constriction to be able to carry out its necessary closure after birth. In humans, patent DA is a life-threatening consequence of prematurity at birth, as oxygenated, high pressure blood is diverted from the aorta back into the left pulmonary artery and thereby into the lungs. A properly closed DA ultimately undergoes physiological fibrosis to evolve into the *ligamentum arteriosum*; untreated, patent DA can lead to congestive heart failure.

Clonal cultures of pigmented, differentiated avian melanocytes have been shown to retain the possibility of either self-renewing or redifferentiating into cells expressing morphological and molecular characteristics of either peripheral glia or pericytes in vitro, even when the original cells were derived from trunk levels [94]. All melanocytes are derived from NCC, in common with some pericytes. Their localization within blood vessels of murine forebrain meninges [93] and the heart [95] in sites shown by chick-quail chimeras to harbor NCC may reflect either a normal but as-yet unidentified secretory function of pigment cells, or simply a sensitized response to signals deployed for pericytic recruitment and vascular stabilization. In favor of the former, leptomeningeal melanocytosis is a hallmark of lethal neurological symptoms in syndromic forms of the rare giant congenital melanocytic nevus [96, 97] and primary pediatric melanoma often occurs in the meninges [98-100]. Consistent with the latter, adult melanoma commonly metastasizes to the same location, all with similarly dire outcomes [101].

Our group has recently observed that in murine models of melanocytosis even devoid of cancer, ectopic melanocytes are also found among the pericytes of capillary beds of the spleen, lymph nodes, gums, adrenal cortex and both male and female genital organs [102]. Remarkably, evidence that vascular pericytes and SMC of mammalian testes can give rise to testosterone-secreting Leydig cells was acquired through a combination of immunohistochemical and proliferation markers, including but not restricted to a neurofilament, GFAP, nestin, NG2, and PDGFRß. However, fate-mapping is still needed to determine the embryonic origins of these pericytes [103].

While there has been no suggestion from examination of quail-chick chimeras that specific testicular cells have a NCC origin, there are differences occasionally observed among distinct vertebrate species or classes. Notable examples are the numerous NCC identified within murine bone marrow stroma [104] and the dental pulp [105], both niches for multipotent stem cell maintenance into adulthood. Such observations raise the hypotheses that pericytic misdifferentiation may play an unexpected role in male infertility, or placental vascular defects [106, 107].

A number of groups have recently turned their efforts to cultivating human pericytes from different sources, either using direct isolation from different tissue sources [108] or in differentiation from stem cell populations of distinct origins. The studies often recombine these cell sources with endothelial cells in order to understand processes of angiogenesis, in effect creating inter-individual chimeric tissues *in vitro* [109], or inter-species chimeras *in vivo* with human-rodent xenografts [110]. Thus, the techniques of experimental embryology appear to be as relevant as ever to understanding pericyte function and their regulation of processes relevant to human health.

